# Minimal cross-over between mutations associated with Omicron variant of SARS-CoV-2 and CD8+ T cell epitopes identified in COVID-19 convalescent individuals

**DOI:** 10.1101/2021.12.06.471446

**Authors:** Andrew D Redd, Alessandra Nardin, Hassen Kared, Evan M Bloch, Brian Abel, Andrew Pekosz, Oliver Laeyendecker, Michael Fehlings, Thomas C Quinn, Aaron AR Tobian

## Abstract

There is a growing concern that ongoing evolution of SARS-CoV-2 could lead to variants of concern (VOC) that are capable of avoiding some or all of the multi-faceted immune response generated by both prior infection or vaccination, with the recently described B.1.1.529 (Omicron) VOC being of particular interest. Peripheral blood mononuclear cell samples from PCR-confirmed, recovered COVID-19 convalescent patients (n=30) infected with SARS-CoV-2 in the United States collected in April and May 2020 who possessed at least one or more of six different HLA haplotypes were selected for examination of their anti-SARS-CoV-2 CD8+ T-cell responses using a multiplexed peptide-MHC tetramer staining approach. This analysis examined if the previously identified viral epitopes targeted by CD8+ T-cells in these individuals (n=52 distinct epitopes) are mutated in the newly described Omicron VOC (n=50 mutations). Within this population, only one low-prevalence epitope from the Spike protein restricted to two HLA alleles and found in 2/30 (7%) individuals contained a single amino acid change associated with the Omicron VOC. These data suggest that virtually all individuals with existing anti-SARS-CoV-2 CD8+ T-cell responses should recognize the Omicron VOC, and that SARS-CoV-2 has not evolved extensive T-cell escape mutations at this time.

**Importance:** The newly identified Omicron variant of concern contains more mutations than any of the previous variants described to date. In addition, many of the mutations associated with the Omicron variant are found in areas that are likely bound by neutralizing antibodies, suggesting that the first line of immunological defense against COVID-19 may be compromised. However, both natural infection and vaccination develop T-cell based responses, in addition to antibodies. This study examined if the parts of the virus, or epitopes, targeted by the CD8+ T-cell response in thirty individuals who recovered from COVID-19 in 2020 were mutated in the Omicron variant. Only one of 52 epitopes identified in this population contained an amino acid that was mutated in Omicron. These data suggest that the T-cell immune response in previously infected, and most likely vaccinated individuals, should still be effective against Omicron.

## Introduction

As the global COVID-19 pandemic enters its third year, there is a growing concern that ongoing evolution of SARS-CoV-2 could lead to a variant of the virus that is capable of avoiding the multi-faceted immune response generated by both prior infection or vaccination[1–3]. Several of these variants of concern (VOC) have been identified throughout the pandemic, and have been associated with large scale waves of infection. To date, while several of these VOC have exhibited varying levels of antibody resistance *in vitro*, vaccination, as well as previous infection by SARS-CoV-2, have been shown to maintain a significant level of protection against breakthrough or re-infections, especially in terms of preventing serious disease and mortality[4,5]. However, the recent description of the B.1.1.529 variant, which was later designated as Omicron, contains a greater number of mutations than the previous VOC[6]. If the mutations in the Omicron VOC mediated resistance from any part of the anti-SARC-CoV-2 immune response, either from vaccination or infection, it would have consequences for efforts to contain the COVID-19 pandemic. The variant has also now been identified on every continent except Antarctica, suggesting it has significant transmission potential, similar to other VOC.

The majority of the mutations associated with the Omicron VOC are located in the Spike protein of the virus, presumably due to selection for evasion of antibody responses, and could have significant effects on the ability of pre-existing antibodies to neutralize the virus, although to what extent this is the case has yet to be determined. It is also unknown how these mutations may affect non-neutralizing binding antibody responses. While it is critical to determine the extent that Omicron may or may not be susceptible to existing humoral responses, T-cell associated immunity is in general significantly more difficult for viruses to overcome due to the broad and adaptable response generated in a given individual, as well as the variety of HLA haplotypes between individuals.

A previous analysis by our group of CD8+ T cell responses to the original SARS-CoV-2 variant in convalescent individuals found a broad and varied immune response in virtually all patients examined, even in individuals with relatively low anti-SARS-CoV-2 antibody responses[1]. A subsequent analysis of these data found that mutations associated with the Alpha, Beta, and Gamma VOC had very minimal cross-over with the epitopes identified in this earlier study (1/52 epitopes affected), suggesting that the CD8+ T cell response from earlier infection would almost certainly still be effective against the new variants[7]. In this study, the mutations associated with Omicron VOC are examined in an identical manner.

## Methods

The detailed methods of the earlier two studies were published previously[1,7]. In short, peripheral blood mononuclear cell (PBMC) samples from PCR-confirmed, recovered COVID-19 convalescent plasma donors collected in April and May 2020 in the Baltimore, MD and Washington DC region who possessed at least one or more of six different HLA haplotypes (HLA-A*01:01, HLA-A*02:01, HLA-A03:01, HLA-A*11:01, HLA-A*24:02 and HLA-B*07:02) were selected for examination of their anti-SARS-CoV-2 CD8+ T cell responses using a multiplexed peptide-MHC tetramer staining approach.

The mutations associated with Omicron VOC (PL: K38R, delta1265, L1266I, A1892T; Nsp4: T492I; 3CL: P132H; Nsp6: delta105-107, I189V; RdRp: P323L; Nsp14:I42V; Spike: A67V, delta69-70, T95I, G142D, delta143-145, N211I; L212V_RE, V213P, R214E, G339D, S371L, S373P, K417N, N440K, G446S, S477N, T478K, E484A, Q493R, G496S, Q498R, T547K, D614G, H655Y, N679K, P681H, N764K, D796Y, N856K, Q954H, N969K, L981F; Envelope: T9I; Matrix: D3G, Q19E, A63T; Nucleocapsid: P13L, delta31-33, R203K, G204R) were mapped to the epitope map developed in the previous analyses, and examined for possible cross-over[8].

### Ethics

All study participants provided written informed consent, and this study was approved by the Johns Hopkins Institutional Review Board.

## Results

The individuals examined in this study were primarily male (60%), and the blood samples were collected a median of 42.5 days (interquartile range 37.5-48.0) from initial diagnosis[1]. The patients were selected from the larger study population according to sample availability, and anti-SARS-CoV-2 IgG responses with ten individuals selected from each of three antibody tertiles[1,3]. 132 SARS-CoV-2-specific CD8+ T cell responses were identified in these individuals, which corresponded to 52 unique epitopes found across the viral genome targeting both structural and non-structural proteins.

Of the mutations associated with the Omicron variant (n=50), only one in the Spike protein (T95I) overlapped with a CD8+ T-cell epitope (GVYFASTEK) identified in this population (Figures 1 and S1). This epitope is restricted to HLA*A03:01 and HLA*A11:01 and a T-cell reactivity was detected in two individuals, typed as HLA:A03:01 and HLA:A03:01/ HLA:A11:01, respectively[9]. Despite the possibility to induce T-cell responses against GVYFASTEK presented on both alleles in one individual, this epitope represented a low-prevalence target in both of these individuals making up 0.1% and a combined 0.4% of all CD8+ T-cell responses in each individual, respectively[1]. In addition, this epitope was 1 of 5 and 1 of 13 of the anti-SARS-CoV-2 epitopes targeted by the two individuals, respectively.

**Figure 1:**
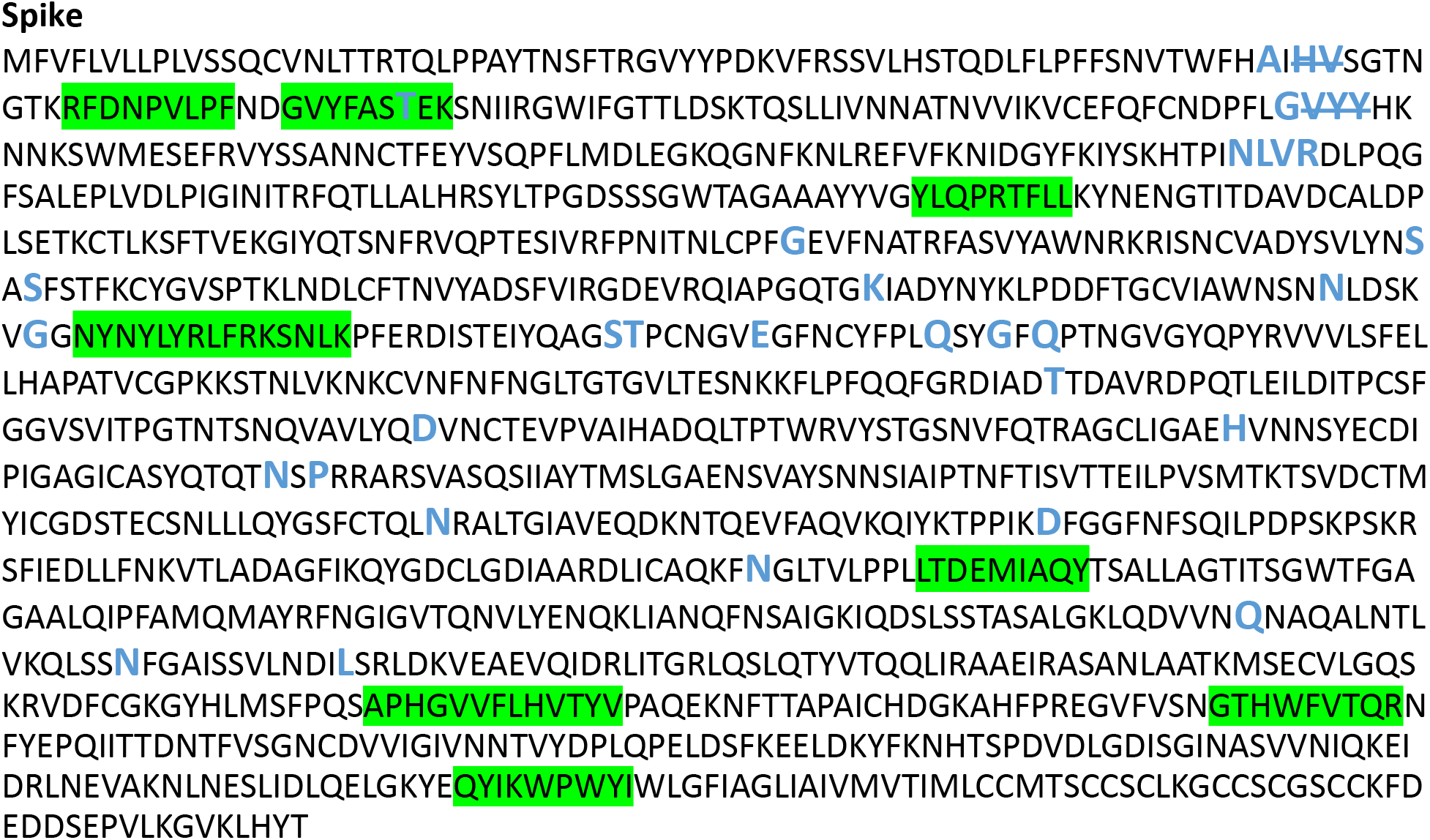
SARS-CoV-2 Wuhan variant protein amino acid sequence for Spike protein with CD8+ T cell epitopes highlighted (green) and all mutations and deletions (slash) sites indicated in large blue letters.

## Discussion

This study demonstrates that despite the substantial number of mutations in the Omicron VOC, in this population only one low-prevalence CD8+ T-cell epitope from the Spike protein contained a single amino acid change. No other mutations were associated with our previously identified epitopes. These data suggest that virtually all individuals with existing anti-SARS-CoV-2 CD8+ T-cell responses should recognize the Omicron VOC, and that SARS-CoV-2 has not evolved extensive T-cell escape mutations at this point.

There are several limitations for this analysis. The results are based on a relatively small samples size of individuals who were all from the United States. Additionally, we examined the CD8+ T-cell responses in previously infected but not vaccinated individuals, and it is possible that the T-cell responses in this later group are more limited and therefore more susceptible to escape. However, work examining T-cell responses from vaccinees has demonstrated a strong CD4+ and CD8+ T-cell responses in these individuals, suggesting that similar trends should be seen in this population as well[2,10].

While it is unknown what specific immune response, or more likely combination of responses, provides optimum protection against SARS-CoV-2 infection and COVID-19, it almost certainly includes a broad and robust CD8+ T-cell response. These data build on the previous analysis of the initial VOC, and confirm that while SARS-CoV-2 has demonstrated a continued pattern of ongoing evolution this has not resulted in any meaningful accumulation of CD8+ T-cell escape mutations[7]. These data also suggest that existing CD8+ T-cell responses from a previous SARS-CoV-2 infection, and most likely from vaccination as well, will still recognize the Omicron VOC and should provide a significant level of protection against COVID-19.

## Acknowledgements

The authors would like to thank the CCP study and laboratory teams, and all the convalescent plasma donors for the generous participation in the study.

## References

1. Kared H, Redd AD, Bloch EM, et al. SARS-CoV-2-specific CD8+ T cell responses in convalescent COVID-19 individuals. J Clin Invest. American Society for Clinical Investigation; 2021; 131(5).

2. Woldemeskel BA, Garliss CC, Blankson JN. SARS-CoV-2 mRNA vaccines induce broad CD4+ T cell responses that recognize SARS-CoV-2 variants and HCoV-NL63. J Clin Invest [Internet]. American Society for Clinical Investigation; 2021 [cited 2021 Nov 30]; 131(10). Available from: https://doi.org/10.1172/JCI149335DS1

3. Klein SL, Pekosz A, Park HS, et al. Sex, age, and hospitalization drive antibody responses in a COVID-19 convalescent plasma donor population. J Clin Invest [Internet]. American Society for Clinical Investigation; 2020 [cited 2021 Feb 5]; 130(11):6141–6150. Available from: https://doi.org/10.1172/JCI142004.

4. Lopez Bernal J, Andrews N, Gower C, et al. Effectiveness of Covid-19 Vaccines against the B.1.617.2 (Delta) Variant. N Engl J Med [Internet]. N Engl J Med; 2021 [cited 2021 Dec 1]; 385(7):585–594. Available from: https://pubmed.ncbi.nlm.nih.gov/34289274/

5. Tang P, Hasan MR, Chemaitelly H, et al. BNT162b2 and mRNA-1273 COVID-19 vaccine effectiveness against the SARS-CoV-2 Delta variant in Qatar. Nat Med [Internet]. Nat Med; 2021 [cited 2021 Dec 1];. Available from: https://pubmed.ncbi.nlm.nih.gov/34728831/

6. WHO. Classification of Omicron (B.1.1.529): SARS-CoV-2 Variant of Concern [Internet]. 2021 [cited 2021 Nov 30]. Available from: https://www.who.int/news/item/26-11-2021-classification-of-omicron-(b.1.1.529)-sars-cov-2-variant-of-concern

7. Redd AD, Nardin A, Kared H, et al. CD8+ T-Cell Responses in COVID-19 Convalescent Individuals Target Conserved Epitopes From Multiple Prominent SARS-CoV-2 Circulating Variants. Open Forum Infect Dis [Internet]. Oxford Academic; 2021 [cited 2021 Aug 25]; 8(7). Available from: https://academic.oup.com/ofid/article/8/7/ofab143/6189113

8. GISAID. GISAID [Internet]. [cited 2021 Dec 1]. Available from: https://www.gisaid.org/

9. Grifoni A, Weiskopf D, Ramirez SI, et al. Targets of T Cell Responses to SARS-CoV-2 Coronavirus in Humans with COVID-19 Disease and Unexposed Individuals. Cell [Internet]. Cell; 2020 [cited 2021 Dec 1]; 181(7):1489–1501.e15. Available from: https://pubmed.ncbi.nlm.nih.gov/32473127/

10. Oberhardt V, Luxenburger H, Kemming J, et al. Rapid and stable mobilization of CD8+ T cells by SARS-CoV-2 mRNA vaccine. Nat 2021 5977875 [Internet]. Nature Publishing Group; 2021 [cited 2021 Nov 30]; 597(7875):268–273. Available from: https://www.nature.com/articles/s41586-021-03841-4

