## Supplemental Figure S1 for "Minimal cross-over between mutations associated with Omicron variant of SARS-CoV-2 and CD8+ T cell epitopes identified in COVID-19 convalescent individuals"

**Figure S1:** SARS-CoV-2 Wuhan variant protein amino acid sequence with CD8+ T cell epitopes highlighted and all mutations and deletions (slash) sites indicated in large blue letters (non-overlapping).

Nucleoprotein

MSDNGPQNQRNA**P**RITFGGPSDSTGSNQNG**~~ERS~~**GARSKQRRPQGLPNNTASWFTALTQHGKEDLKFPRGQGVPINTNSSPDDQIGYYRRATRRIRGGDGKMKDLSPRWYFYYLGTGPEAGLPYGANKDGIIWVATEGALNTPKDHIGTRNPANNAAIVLQLPQGTTLPKGFYAEGSRGGSQASSRSSSRSRNSSRNSTPGSS**RG**TSPARMAGNGGDAALALLLLDRLNQLESKMSGKGQQQQGQTVTKKSAAEASKKPRQKRTATKAYNVTQAFGRRGPEQTQGNFGDQELIRQGTDYKHWPQIAQFAPSASAFFGMSRIGMEVTPSGTWLTYTGAIKLDDKDPNFKDQVILLNKHIDAYKTFPPTEPKKDKKKKADETQALPQRQKKQQTVTLLPAADLDDFSKQLQQSMSSADSTQA

ORF7a

MKIILFLALITLATCELYHYQECVRGTTVLLKEPCSSGTYEGNSPFHPLADNKFALTCFSTQFAFACPDGVKHVYQLRARSVSPKLFIRQEEVQELYSPIFLIVAAIVFITLCFTLKRKTE

ORF3a

MDLFMRIFTIGTVTLKQGEIKDATPSDFVRATATIPIQASLPFGWLIVGVALLAVFQSASKIITLKKRWQLALSKGVHFVCNLLLLFVTVYSHLLLVAAGLEAPFLYLYALVYFLQSINFVRIIMRLWLCWKCRSKNPLLYDANYFLCWHTNCYDYCIPYNSVTSSIVITSGDGTTSPISEHDYQIGGYTEKWESGVKDCVVLHSYFTSDYYQLYSTQLSTDTGVEHVTFFIYNKIVDEPEEHVQIHTIDGSSGVVNPVMEPIYDEPTTTTSVPL

**ORF1ab**

Nsp1/2

MESLVPGFNEKTHVQLSLPVLQVRDVLVRGFGDSVEEVLSEARQHLKDGTCGLVEVEKGVLPQLEQPYVFIKRSDARTAPHGHVMVELVAELEGIQYGRSGETLGVLVPHVGEIPVAYRKVLLRKNGNKGAGGHSYGADLKSFDLGDELGTDPYEDFQENWNTKHSSGVTRELMRELNGGAYTRYVDNNFCGPDGYPLECIKDLLARAGKASCTLSEQLDFIDTKRGVYCCREHEHEIAWYTERSEKSYELQTPFEIKLAKKFDTFNGECPNFVFPLNSIIKTIQPRVEKKKLDGFMGRIRSVYPVASPNECNQMCLSTLMKCDHCGETSWQTGDFVKATCEFCGTENLTKEGATTCGYLPQNAVVKIYCPACHNSEVGPEHSLAEYHNESGLKTILRKGGRTIAFGGCVFSYVGCHNKCAYWVPRASANIGCNHTGVVGEGSEGLNDNLLEILQKEKVNINIVGDFKLNEEIAIILASFSASTSAFVETVKGLDYKAFKQIVESCGNFKVTKGKAKKGAWNIGEQKSILSPLYAFASEAARVVRSIFSRTLETAQNSVRVLQKAAITILDGISQYSLRLIDAMMFTSDLATNNLVVMAYITGGVVQLTSQWLTNIFGTVYEKLKPVLDWLEEKFKEGVEFLRDGWEIVKFISTCACEIVGGQIVTCAKEIKESVQTFFKLVNKFLALCADSIIIGGAKLKALNLGETFVTHSKGLYRKCVKSREETGLLMPLKAPKEIIFLEGETLPTEVLTEEVVLKTGDLQPLEQPTSEAVEAPLVGTPVCINGLMLLEIKDTEKYCALAPNMMVTNNTFTLKGG

Nsp3

APTKVTFGDDTVIEVQGYKSVNITFELDERIDKVLNE**K**CSAYTVELGTEVNEFACVVADAVIKTLQPVSELLTPLGIDLDEWSMATYYLFDESGEFKLASHMYCSFYPPDEDEEEGDCEEEEFEPSTQYEYGTEDDYQGKPLEFGATSAALQPEEEQEEDWLDDDSQQTVGQQDGSEDNQTTTIQTIVEVQPQLEMELTPVVQTIEVNSFSGYLKLTDNVYIKNADIVEEAKKVKPTVVVNAANVYLKHGGGVAGALNKATNNAMQVESDDYIATNGPLKVGGSCVLSGHNLAKHCLHVVGPNVNKGEDIQLLKSAYENFNQHEVLLAPLLSAGIFGADPIHSLRVCVDTVRTNVYLAVFDKNLYDKLVSSFLEMKSEKQVEQKIAEIPKEEVKPFITESKPSVEQRKQDDKKIKACVEEVTTTLEETKFLTENLLLYIDINGNLHPDSATLVSDIDITFLKKDAPYIVGDVVQEGVLTAVVIPTKKAGGTTEMLAKALRKVPTDNYITTYPGQGLNGYTVEEAKTVLKKCKSAFYILPSIISNEKQEILGTVSWNLREMLAHAEETRKLMPVCVETKAIVSTIQRKYKGIKIQEGVVDYGARFYFYTSKTTVASLINTLNDLNETLVTMPLGYVTHGLNLEEAARYMRSLKVPATVSVSSPDAVTAYNGYLTSSSKTPEEHFIETISLAGSYKDWSYSGQSTQLGIEFLKRGDKSVYYTSNPTTFHLDGEVITFDNLKTLLSLREVRTIKVFTTVDNINLHTQVVDMSMTYGQQFGPTYLDGADVTKIKPHNSHEGKTFYVLPNDDTLRVEAFEYYHTTDPSFLGRYMSALNHTKKWKYPQVNGLTSIKWADNNCYLATALLTLQQIELKFNPPALQDAYYRARAGEAANFCALILAYCNKTVGELGDVRETMSYLFQHANLDSCKRVLNVVCKTCGQQQTTLKGVEAVMYMGTLSYEQFKKGVQIPCTCGKQATKYLVQQESPFVMMSAPPAQYELKHGTFTCASEYTGNYQCGHYKHITSKETLYCIDGALLTKSSEYKGPITDVFYKENSYTTTIKPVTYKLDG**V**VCTEIDPKLDNYYKKDNSYFTEQPIDLVPNQPYPNASFDNFKFVCDNIKFADDLNQLTGYKKPASRELKVTFFPDLNGDVVAIDYKHYTPSFKKGAKLLHKPIVWHVNNATNKATYKPNTWCIRCLWSTKPVETSNSFDVLKSEDAQGMDNLACEDLKPVSEEVVENPTIQKDVLECNVKTTEVVGDIILKPANN**~~S~~L**KITEEVGHTDLMAAYVDNSSLTIKKPNELSRVLGLKTLATHGLAAVNSVPWDTIANYAKPFLNKVVSTTTNIVTRCLNRVCTNYMPYFFTLLLQLCTFTRSTNSRIKASMPTTIAKNTVKSVGKFCLEASFNYLKSPNFSKLINIIIWFLLLSVCLGSLIYSTAALGVLMSNLGMPSYCTGYREGYLNSTNVTIATYCTGSIPCSVCLSGLDSLDTYPSLETIQITISSFKWDLTAFGLVAEWFLAYILFTRFFYVLGLAAIMQLFFSYFAVHFISNSWLMWLIINLVQMAPISAMVRMYIFFASFYYVWKSYVHVVDGCNSSTCMMCYKRNRATRVECTTIVNGVRRSFYVYANGGKGFCKLHNWNCVNCDTFCAGSTFISDEVARDLSLQFKRPINPTDQSSYIVDSVTVKNGSIHLYFDKAGQKTYERHSLSHFVNLDNLRANNTKGSLPINVIVFDGKSKCEESSAKSASVYYSQLMCQPILLLDQALVSDVGDSAEVAVKMFDAYVNTFSSTFNVPMEKLKTLVATAEAELAKNVSLDNVLSTFISAARQGFVDSDVETKDVVECLKLSHQSDIEVTGDSCNNYMLTYNKVENMTPRDLGACIDCSARHINAQVAKSHNI**A**LIWNVKDFMSLSEQLRKQIRSAAKKNNLPFKLTCATTRQVVNVVTTKIALKGG

Nsp4

KIVNNWLKQLIKVTLVFLFVAAIFYLITPVHVMSKHTDFSSEIIGYKAIDGGVTRDIASTDTCFANKHADFDTWFSQRGGSYTNDKACPLIAAVITREVGFVVPGLPGTILRTTNGDFLHFLPRVFSAVGNICYTPSKLIEYTDFATSACVLAAECTIFKDASGKPVPYCYDTNVLEGSVAYESLRPDTRYVLMDGSIIQFPNTYLEGSVRVVTTFDSEYCRHGTCERSEAGVCVSTSGRWVLNNDYYRSLPGVFCGVDAVNLLTNMFTPLIQPIGALDISASIVAGGIVAIVVTCLAYYFMRFRRAFGEYSHVVAFNTLLFLMSFTVLCLTPVYSFLPGVYSVIYLYLTFYLTNDVSFLAHIQWMVMFTPLVPFWITIAYIICISTKHFYWFFSNYLKRRVVFNGVSFSTFEEAALCTFLLNKEMYLKLRSDVLLPLTQYNRYLALYNKYKYFSGAMDTTSYREAACCHLAKALNDFSNSGSDVLYQPPQ**T**SITSAVLQ

Nsp5

SGFRKMAFPSGKVEGCMVQVTCGTTTLNGLWLDDVVYCPRHVICTSEDMLNPNYEDLLIRKSNHNFLVQAGNVQLRVIGHSMQNCVLKLKVDTANPKTPKYKFVRIQPGQTFSVLACYNGSPSGVYQCAMR**P**NFTIKGSFLNGSCGSVGFNIDYDCVSFCYMHHMELPTGVHAGTDLEGNFYGPFVDRQTAQAAGTDTTITVNVLAWLYAAVINGDRWFLNRFTTTLNDFNLVAMKYNYEPLTQDHVDILGPLSAQTGIAVLDMCASLKELLQNGMNGRTILGSALLEDEFTPFDVVRQCSGVTFQ

Nsp6

SAVKRTIKGTHHWLLLTILTSLLVLVQSTQWSLFFFLYENAFLPFAMGIIAMSAFAMMFVKHKHAFLCLFLLPSLATVAYFNMVYMPASWVMRIMTWLDMVDTS**~~LSG~~**FKLKDCVMYASAVVLLILMTARTVYDDGARRVWTLMNVLTLVYKVYYGNALDQAISMWALIISVTSNYSGVVTTVMFLARG**I**VFMCVEYCPIFFITGNTLQCIMLVYCFLGYFCTCYFGLFCLLNRYFRLTLGVYDYLVSTQEFRYMNSQGLLPPKNSIDAFKLNIKLLGVGGKPCIKVATVQ

Nsp7-11

SKMSDVKCTSVVLLSVLQQLRVESSSKLWAQCVQLHNDILLAKDTTEAFEKMVSLLSVLLSMQGAVDINKLCEEMLDNRATLQAIASEFSSLPSYAAFATAQEAYEQAVANGDSEVVLKKLKKSLNVAKSEFDRDAAMQRKLEKMADQAMTQMYKQARSEDKRAKVTSAMQTMLFTMLRKLDNDALNNIINNARDGCVPLNIIPLTTAAKLMVVIPDYNTYKNTCDGTTFTYASALWEIQQVVDADSKIVQLSEISMDNSPNLAWPLIVTALRANSAVKLQNNELSPVALRQMSCAAGTTQTACTDDNALAYYNTTKGGRFVLALLSDLQDLKWARFPKSDGTGTIYTELEPPCRFVTDTPKGPKVKYLYFIKGLNNLNRGMVLGSLAATVRLQAGNATEVPANSTVLSFCAFAVDAAKAYKDYLASGGQPITNCVKMLCTHTGTGQAITVTPEANMDQESFGGASCCLYCRCHIDHPNPKGFCDLKGKYVQIPTTCANDPVGFTLKNTVCTVCGMWKGYGCSCDQLREPMLQ

Nsp12/polymerase

SADAQSFLNRVCGVSAARLTPCGTGTSTDVVYRAFDIYNDKVAGFAKFLKTNCCRFQEKDEDDNLIDSYFVVKRHTFSNYQHEETIYNLLKDCPAVAKHDFFKFRIDGDMVPHISRQRLTKYTMADLVYALRHFDEGNCDTLKEILVTYNCCDDDYFNKKDWYDFVENPDILRVYANLGERVRQALLKTVQFCDAMRNAGIVGVLTLDNQDLNGNWYDFGDFIQTTPGSGVPVVDSYYSLLMPILTLTRALTAESHVDTDLTKPYIKWDLLKYDFTEERLKLFDRYFKYWDQTYHPNCVNCLDDRCILHCANFVLFSTVFP**P**TSFGPLVRKIFVDGVPFVVSTGYHFRELGVVHNQDVNLHSSRLSFKELLVYAADPAMHAASGNLLLDKRTTCFSVAALTNNVAFQTVKPGNFNKDFYDFAVSKGFFKEGSSVELKHFFFAQDGNAAISDYDYYRYNLPTMCDIRQLLFVVEVVDKYFDCYDGGCINANQVIVNNLDKSAGFPFNKWGKARLYYDSMSYEDQDALFAYTKRNVIPTITQMNLKYAISAKNRARTVAGVSICSTMTNRQFHQKLLKSIAATRGATVVIGTSKFYGGWHNMLKTVYSDVENPHLMGWDYPKCDRAMPNMLRIMASLVLARKHTTCCSLSHRFYRLANECAQVLSEMVMCGGSLYVKPGGTSSGDATTAYANSVFNICQAVTANVNALLSTDGNKIADKYVRNLQHRLYECLYRNRDVDTDFVNEFYAYLRKHFSMMILSDDAVVCFNSTYASQGLVASIKNFKSVLYYQNNVFMSEAKCWTETDLTKGPHEFCSQHTMLVKQGDDYVYLPYPDPSRILGAGCFVDDIVKTDGTLMIERFVSLAIDAYPLTKHPNQEYADVFHLYLQYIRKLHDELTGHMLDMYSVMLTNDNTSRYWEPEFYEAMYTPHTVLQ

Nsp13

AVGACVLCNSQTSLRCGACIRRPFLCCKCCYDHVISTSHKLVLSVNPYVCNAPGCDVTDVTQLYLGGMSYYCKSHKPPISFPLCANGQVFGLYKNTCVGSDNVTDFNAIATCDWTNAGDYILANTCTERLKLFAAETLKATEETFKLSYGIATVREVLSDRELHLSWEVGKPRPPLNRNYVFTGYRVTKNSKVQIGEYTFEKGDYGDAVVYRGTTTYKLNVGDYFVLTSHTVMPLSAPTLVPQEHYVRITGLYPTLNISDEFSSNVANYQKVGMQKYSTLQGPPGTGKSHFAIGLALYYPSARIVYTACSHAAVDALCEKALKYLPIDKCSRIIPARARVECFDKFKVNSTLEQYVFCTVNALPETTADIVVFDEISMATNYDLSVVNARLRAKHYVYIGDPAQLPAPRTLLTKGTLEPEYFNSVCRLMKTIGPDMFLGTCRRCPAEIVDTVSALVYDNKLKAHKDKSAQCFKMFYKGVITHDVSSAINRPQIGVVREFLTRNPAWRKAVFISPYNSQNAVASKILGLPTQTVDSSQGSEYDYVIFTQTTETAHSCNVNRFNVAITRAKVGILCIMSDRDLYDKLQFTSLEIPRRNVATLQ

Nsp14

AENVTGLFKDCSKVITGLHPTQAPTHLSVDTKFKTEGLCVD**I**PGIPKDMTYRRLISMMGFKMNYQVNGYPNMFITREEAIRHVRAWIGFDVEGCHATREAVGTNLPLQLGFSTGVNLVAVPTGYVDTPNNTDFSRVSAKPPPGDQFKHLIPLMYKGLPWNVVRIKIVQMLSDTLKNLSDRVVFVLWAHGFELTSMKYFVKIGPERTCCLCDRRATCFSTASDTYACWHHSIGFDYVYNPFMIDVQQWGFTGNLQSNHDLYCQVHGNAHVASCDAIMTRCLAVHECFVKRVDWTIEYPIIGDELKINAACRKVQHMVVKAALLADKFPVLHDIGNPKAIKCVPQADVEWKFYDAQPCSDKAYKIEELFYSYATHSDKFTDGVCLFWNCNVDRYPANSIVCRFDTRVLSNLNLPGCDGGSLYVNKHAFHTPAFDKSAFVNLKQLPFFYYSDSPCESHGKQVVSDIDYVPLKSATCITRCNLGGAVCRHHANEYRLYLDAYNMMISAGFSLWVYKQFDTYNLWNTFTRLQ

Nsp15/16

SLENVAFNVVNKGHFDGQQGEVPVSIINNTVYTKVDGVDVELFENKTTLPVNVAFELWAKRNIKPVPEVKILNNLGVDIAANTVIWDYKRDAPAHISTIGVCSMTDIAKKPTETICAPLTVFFDGRVDGQVDLFRNARNGVLITEGSVKGLQPSVGPKQASLNGVTLIGEAVKTQFNYYKKVDGVVQQLPETYFTQSRNLQEFKPRSQMEIDFLELAMDEFIERYKLEGYAFEHIVYGDFSHSQLGGLHLLIGLAKRFKESPFELEDFIPMDSTVKNYFITDAQTGSSKCVCSVIDLLLDDFVEIIKSQDLSVVSKVVKVTIDYTEISFMLWCKDGHVETFYPKLQSSQAWQPGVAMPNLYKMQRMLLEKCDLQNYGDSATLPKGIMMNVAKYTQLCQYLNTLTLAVPYNMRVIHFGAGSDKGVAPGTAVLRQWLPTGTLLVDSDLNDFVSDADSTLIGDCATVHTANKWDLIISDMYDPKTKNVTKENDSKEGFFTYICGFIQQKLALGGSVAIKITEHSWNADLYKLMGHFAWWTAFVTNVNASSSEAFLIGCNYLGKPREQIDGYVMHANYIFWRNTNPIQLSSYSLFDMSKFPLKLRGTAVMSLKEGQINDMILSLLSKGRLIIRENNRVVISSDVLVNN
